# Diverging asymmetry of intrinsic functional organization in autism

**DOI:** 10.1101/2023.04.05.535683

**Authors:** Bin Wan, Seok-Jun Hong, Richard A.I. Bethlehem, Dorothea L. Floris, Boris C. Bernhardt, Sofie L. Valk

## Abstract

Autism is a neurodevelopmental condition involving atypical sensory-perceptual functions together with language and socio-cognitive deficits. Previous work has reported subtle alterations in the asymmetry of brain structure and reduced laterality of functional activation in individuals with autism relative to non-autistic individuals (NAI). However, whether functional asymmetries show altered intrinsic systematic organization in autism remains unclear. Here, we computed inter- and intra-hemispheric asymmetry of intrinsic functional gradients capturing connectome organization along three axes, stretching between sensory-default, somatomotor-visual, and default-multiple demand networks, to study system-level hemispheric imbalances in autism. We observed decreased leftward functional asymmetry of language network organization in individuals with autism, relative to NAI. Whereas language network asymmetry varied across age groups in NAI, this was not the case in autism, suggesting atypical functional laterality in autism may result from altered developmental trajectories. Finally, we observed that intra-but not inter-hemispheric features were predictive of the severity of autistic traits. In sum, our findings illustrate how regional and patterned functional lateralization is altered in autism at the system level. Such differences may be rooted in altered developmental trajectories of functional organization asymmetry in autism.

## Introduction

Autism is a heterogeneous neurodevelopmental condition with a prevalence exceeding 2% in a recent survey in the U.S. (Xu et al., 2018). It is characterized by life-long differences in social interaction and communication alongside restricted and repetitive interests/behaviors (American Psychiatric Association, 2013). The widespread behavioral differences observed in individuals with autism are paralleled by reports of structural and functional alterations in both sensory and association regions of the brain (Chen et al., 2021; Hong et al., 2019; Jung et al., 2014; Nunes et al., 2019; Oldehinkel et al., 2019; Ornitz, 1974; B. Park et al., 2021; S. Park et al., 2021; Valk et al., 2015; Washington et al., 2014).

Whole brain differences in structure and function between autism and non-autistic individuals (NAI) are augmented by observations of disrupted patterns of brain asymmetry (Postema et al., 2019; Sha et al., 2022), possibly linked to abnormal lateralisation of functional processes supporting language and social cognition (Floris et al., 2021; Floris & Howells, 2018; Jouravlev et al., 2020; Just et al., 2004; Lindell & Hudry, 2013; Nielsen et al., 2014). Asymmetry is a key feature of brain organization, supporting a flexible interplay between specific local neural modules linked to functional specialization underlying human cognition (Hartwigsen et al., 2021). In particular, left-hemispheric regions are biased to interact stronger within the hemisphere, whereas interactions of the right hemispheric regions are more balanced (Gotts et al., 2013). Recent work has shown that individuals with autism display marked and widespread atypical patterns of asymmetry of local structure (Postema et al., 2019). Such differences may reflect changes in network-level embedding, in particular in association regions, as measured by structural covariance (Sha et al., 2022). Functionally, individuals with autism exhibit idiosyncratic alterations in homotopic inter-hemispheric connectivity patterns, indicating more variation in the autistic population (Benkarim et al., 2021; Hahamy et al., 2015). Further, they show atypical rightward functional lateralisation in mean motor circuit connectivity (Floris et al., 2016). Independent component analysis (ICA) using the resting state functional connectome suggests that component loadings are more rightward in individuals with autism (Cardinale et al., 2013). Last, decreased asymmetry of functional activation patterning has been observed in individuals with autism during the letter fluency task (Kleinhans et al., 2008). Functional differences may be rooted in altered developmental trajectories of functional lateralization in autism (Flagg et al., 2005), leading to altered global features of brain organization and asymmetry, as captured by low dimensional connectome embeddings (Hong et al., 2020; Vos de Wael et al., 2020). Thus, observed localized asymmetry differences may reflect altered system-wide functional organization.

To further understand system-level functional lateralization alterations in autism, here we investigated the asymmetry of intra- and inter-hemispheric functional connectome gradients (Gotts et al., 2013; Wan et al., 2022), which robustly capture associated organizational features (Margulies et al., 2016; Vos de Wael et al., 2020). Cortical regions show organizational axes reflecting integration and segregation (Gotts et al., 2013) along different dimensions (Huntenburg et al., 2018; Margulies et al., 2016; Paquola et al., 2022; Smallwood et al., 2021). Gradient 1 (G1) transitions between sensory and default mode networks, gradient 2 (G2) between sensory and visual cortices, and gradient 3 (G3) between default mode and multiple demand networks. These gradients can be reliably identified (Hong et al., 2020), and are among the most widely studied in the literature (Bernhardt et al., 2022). Together, these gradients describe patterns of developmental and heritable variation in the human cortex (Dong et al., 2021; Valk et al., 2022; Xia et al., 2022). In previous work, we and others have shown robust hemispheric differences in functional organization in non-autistic adults (Gonzalez Alam et al., 2021; Liang et al., 2021; Wan et al., 2022). For example, the main lateralization axis separated different abstract cognitive functions (language to executive function) from left to right (Wan et al., 2022). Given that language impairments and verbal imbalances are key traits of autism (Boucher, 2012; Hong et al., 2023; Kjellmer et al., 2018; Vogindroukas et al., 2022) and executive function may underlie the psychological and behavioral neurodivergence observed in autism (Demetriou et al., 2018; Zhang et al., 2020), we hypothesize that atypical lateralization axes in autism may contribute to autistic behaviors.

To answer our research question, we first compared the asymmetry of functional gradients between autistic individuals and NAI to reveal the differences between groups. Because brain asymmetry (Kong et al., 2018; Roe et al., 2021) and gradients (Bethlehem et al., 2020; Dong et al., 2021; Xia et al., 2022) are affected by age, we also evaluated the interaction of age and autism status to reveal the cross-sectional developmental trajectory. Given the heritability of functional gradient asymmetry (Wan et al., 2022), and also of autism (Colvert et al., 2015; Tick et al., 2016), we used prior heritability estimates (Wan et al., 2022), to evaluate whether autism is associated with differences in regions found to be heritable in adulthood. Finally, supervised machine learning was used to establish phenotypical relevance. We also tested robustness using the functional connectome after global signal regression (GSR).

## Results

### Data demographics

We utilized resting-state fMRI data from 5 sites from the Autism Brain Imaging Data Exchange (ABIDE-I) (Di Martino et al., 2014) including: New York University Langone Medical Center (NYU-I, n = 86), University of Pittsburgh, School of Medicine (Pitt, n = 39), and University of Utah, School of Medicine (USM, n = 83), as well as Trinity Centre for Health Sciences, Trinity College Dublin (TCD, n = 32) and NYU-II (n = 43) from ABIDE-II (Di Martino et al., 2017). We selected those sites that included children, adolescents, and adults. All participants were male (n _autism_ = 140 and n _NTC_ = 143) with age ranging from 5 - 40 years. There was no significant age difference (*t* = -0.030, *p* > 0.05) between individuals with autism and NAI. The resting state fMRI data were preprocessed based on C-PAC (https://fcp-indi.github.io/). Functional connectome gradients were computed by aligning each individual to the group-level gradients template with Procrustes rotations by the Human Connectome Project (HCP) (Wan et al., 2022): sensory-default (G1), somatomotor-visual (G2), and default-multiple demand (G3) gradients. The inclusion and exclusion criteria and detailed computation can be seen in the **Methods**.

The full intelligence quotient (FIQ) and Autism Diagnostic Observation Schedule (ADOS, Generic version) score are shown in **Supplementary Table S1**. Of note, there are differences between autism and NAI in FIQ (*t* = -5.710, *p* < 0.001) and head motion (*t* = 2.636, *p* = 0.009). Multi-site effect was removed before analyses via data harmonization that follows an empirical Bayesian approach to balance the effects of each scanner/batch (Fortin et al., 2018).

### Asymmetry along functional organization axes (Figure 1)

We first computed the functional connectome for each individual, and applied diffusion embedding (Margulies et al., 2016; Vos de Wael et al., 2020) to decompose the first 10 gradients of different connectivity patterns (i.e., LL: left-left, LR: left-right, RL: right-left, and RR: right-right). Then, we aligned individual gradients of all the participants to the HCP group-level gradient of the left-left functional connectivity pattern (Wan et al., 2022) with Procrustes rotations. This allowed direct comparison of the organization of functional asymmetry across groups and individuals, in line with previous work (Hong et al., 2019; B. Park et al., 2021; S. Park et al., 2021; Wan et al., 2022). Individual functional gradient computation and analyses with Python packages BrainSpace (Vos de Wael et al., 2020) and BrainStat (Larivière et al., 2023) are described in the **Methods**.

**Figure 1.**
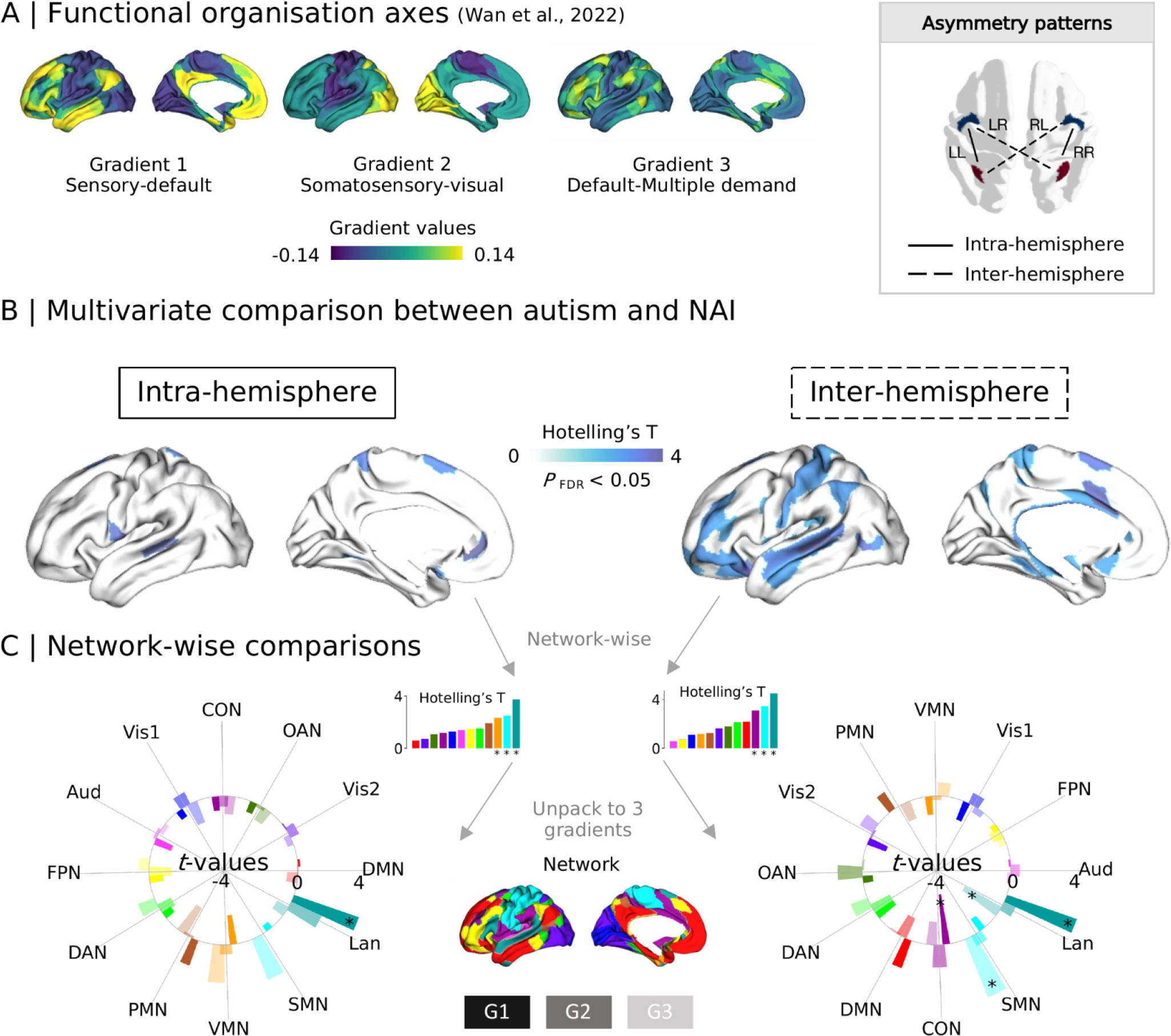
Comparison between individuals with autism and NAI in gradient asymmetry index along functional organizational axes. **A** | HCP group-level gradients of left-left (LL) functional connectome described by (Wan et al., 2022) including G1: sensory-default gradient, G2: somatosensory-visual gradient, and G3: default-multiple demand gradient. Asymmetry has intra- and inter-hemispheric patterns derived from [LL, RR] and [LR, RL] functional connectome. **B** | Multivariate comparison across G1, G2, and G3. The brain maps show the Hotelling’s T values (*p*_FDR_ < 0.05) for multivariate comparison. **C** | Network-wise comparisons. The multivariate analyses were summarized from parcel-wise comparison using multi-modal parcellation (Glasser et al., 2016) to network-wise comparison using Cole-Anticevic (CA) atlas (Ji et al., 2019). Radar-bar plots show network-wise decomposition results (from multivariate to single gradient). Dark, middle dark, and light colors indicate *t*-values of each network along G1, G2, and G3. * marks significant networks. NAI: non-autistic individuals, Vis1: primary visual network, Vis2: secondary visual network, SMN: somatomotor network, CON: cingulo-opercular network, DAN: dorsal attention network, Lan.:language network, FPN: frontoparietal network, Aud.: auditory network, DMN: default mode network, PMN: posterior multimodal network, VMN: ventral multimodal network, OAN: orbito-affective network.

Next, we calculated the asymmetry index (AI) along the three organizational axes (**Figure 1A**) for intra-hemispheric FC patterns (LL minus RR) and inter-hemispheric FC patterns (LR minus RL) following previous work (Wan et al., 2022). Overall, the spatial asymmetric pattern was similar to the HCP asymmetric pattern (Wan et al., 2022), with NAI showing more similar patterns than autism (**Supplementary results)**. We then took a multivariate approach using Hotelling’s *T* to discover shared effects across the three eigenvectors. In *post-hoc* analyses we further investigated contributions of individual gradients to the overall effects, correcting for the number of gradients considered (*p* < 0.05/3). For this analysis, age effect was entered as a covariate during data harmonization.

Parcel-wise multivariate analyses with *p*_FDR_ < 0.05 mapped overall differences between individuals with autism and NAI (**Figure 1B**). This revealed group differences in language-related and somatosensory areas for intra-hemispheric patterns, and inter-hemispheric differences in dorsal prefrontal, superior temporal, and postcentral cortices. We performed *post-hoc* single gradient comparisons of these parcels (**Supplementary results and Table S2**). Positive and negative *t*-values indicate lower and higher left-right asymmetry in individuals with autism relative to NAI. In particular, for intra-hemispheric G1, parcels included medial posterior superior frontal lobule (SFL, *t* = 2.758, *p* = 0.006), area 43 (posterior opercular, *t* = -3.058, *p* = 0.002), and the dorsal posterior superior temporal sulcus (STSdp, *t* = 3.796, *p* < 0.001). For inter-hemispheric G1 parcels included area 33pr (anterior cingulate, *t* = -2.436, *p* = 0.015), area a24pr (anterior cingulate, *t* = -4.390, *p* < 0.001), area p32pr (anterior cingulate, *t* = -3.548, *p* < 0.001), area 47m (frontal pole, *t* = 2.648, *p* = 0.009), area 47s (frontal pole, *t* = 3.384, *p* < 0.001), auditory 5 complex (A5, *t* = 2.813, *p* = 0.005), dorsal anterior superior temporal sulcus (STSda, *t* = 3.838, *p* < 0.001), and temporo-parieto-occipital junction 1 (TPOJ1, *t* = 2.815, *p* = 0.005). The parcel labels refer to Glasser et al., (2016). G2 and G3 showed less strong asymmetric differences between individuals with autism and NAI and have been described in the **Supplementary Results**.

When evaluating network-wise asymmetries (Ji et al., 2019), we observed four significant networks for multivariate comparisons after FDR correction (**Figure 1B** and **Supplementary Table S3**). Only three were observed with statistical significance in single-gradient analyses (**Figure 1C** and **Supplementary Table S3**). Specifically, the language network (Lan., intra-hemispheric G1, *t* = 3.682, *p* < 0.001; inter-hemispheric G1, *t* = 3.973, *p* < 0.001), cingulo-opercular network (CON, inter-hemispheric G1, *t* = -2.248, *p* = 0.007), and somatomotor network (SMN, inter-hemispheric G3, *t* = 3.443, *p* < 0.001) showed differentiable asymmetry. Results remained robust when performing GSR. Detailed reports can be found in **Supplementary Figure S2.** Findings did not change after including FIQ and head motion as covariates during data harmonization.

Inter-subject similarity analyses across each data site (Hahamy et al., 2015) tested the populational differences between individuals with autism and NAI in cortical functional asymmetric patterns. We observed that individuals with autism showed a lower similarity score relative to NAI along the three axes. Detailed reports are shown in **Supplementary Results and Table S8**. This suggests that autism is quite heterogeneous in terms of functional organization asymmetry.

### Developmental effects (Figure 2)

To explore whether the asymmetry of functional gradients develops differently between individuals with autism and NAI, we categorized participants into three age groups including children (5 - 12 years, n = 74), adolescents (12 - 18 years, n = 93), and adults (18 - 40 years, n = 130).

We first examined whether there were age differences within autism and NAI groups. In the comparisons between age groups, we set *p* < 0.05/3 (Bonferroni correction) as the significance level. We observed no significant asymmetry changes with age in autism. However, there were significant age differences in Vis1, Lan., and OAN in NAI. For example, in NAI, children showed increased leftward asymmetry relative to adults in Lan. along G3 (intra-hemispheric, *t* = -3.852, *p* < 0.001; inter-hemispheric, *t* = -2.443, *p* = 0.016). See **Supplementary Results and Table S4** for further details. We then studied the interaction between age and autism status to evaluate whether the age effects are different between autism and NAI. Parcel-wise multivariate analyses revealed interaction effects of age with autism status in parcels primarily located in dorsolateral prefrontal and posterior temporal cortices for the intra-hemispheric pattern, and in parcels mainly located in postcentral and visual cortices for the inter-hemispheric pattern (**Figure 2A**). The detailed parcel-wise and single-gradient results are presented in **Supplementary Table S5**. Regarding network-wise comparisons, **Figure 2B** illustrates intra- and inter-hemispheric patterns of age by autism status effects (**Supplementary Table S5**). Among them, we found interaction effects in Lan. along intra-hemispheric G3 (*t* = 3.830, *p* < 0.001) after Bonferroni correction. Interaction between autism status and age using GSR replicated the intra-hemispheric asymmetry results but not the inter-hemispheric asymmetry results (**Supplementary Results**). Results did not change after including FIQ and head motion as covariates during data harmonization.

**Figure 2.**
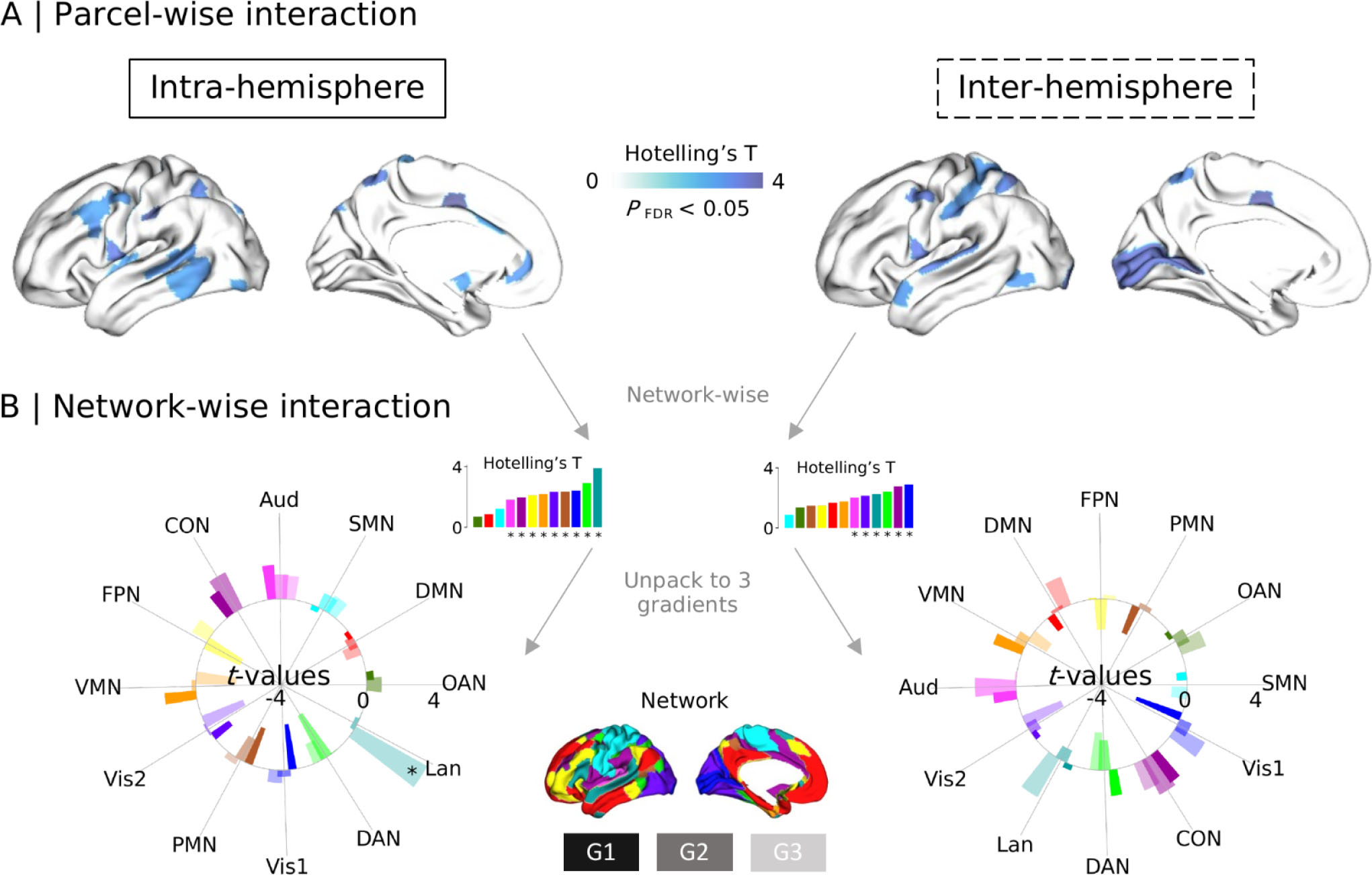
Interaction effects between age and autism status on asymmetry index (AI). **A** | Parcel-wise interaction using multivariate analyses. The brain maps show the Hotelling’s T values (*p*_FDR_ < 0.05) for multivariate comparison across the three gradients. **B** | Network-wise interaction from multivariate to single gradient. Vis1: primary visual network, Vis2: secondary visual network, SMN: somatomotor network, CON: cingulo-opercular network, DAN: dorsal attention network, Lan.:language network, FPN: frontoparietal network, Aud.: auditory network, DMN: default mode network, PMN: posterior multimodal network, VMN: ventral multimodal network, OAN: orbito-affective network.

Analyses for diagnostic differences in each age group have been shown in **Supplementary Results and Table S6**. Overall, diagnostic differences in Lan. along G1 and SMN along G3 were present in adolescents but not in children or adults.

### Meta-analytic functional decoding and heritability (Figure 3)

Having established marked alterations in asymmetry of functional organization between individuals with autism and NAI, which varied across age-groups, we further aimed to contextualize the findings. In particular, to explore how the differences between autism and NAI are related to cognitive functions, we performed meta-analytic decoding using NeuroSynth (Yarkoni et al., 2011) using 24 terms-related z-activation maps, similar to previous work (Margulies et al., 2016; Valk et al., 2022; Wan et al., 2022). Second, we performed decoding of asymmetry effects relative to heritability of asymmetry observed in previous work, based on HCP twin-based data (**Figure 3A**) from (Wan et al., 2022). Details can be found in the **Methods**.

**Figure 3.**
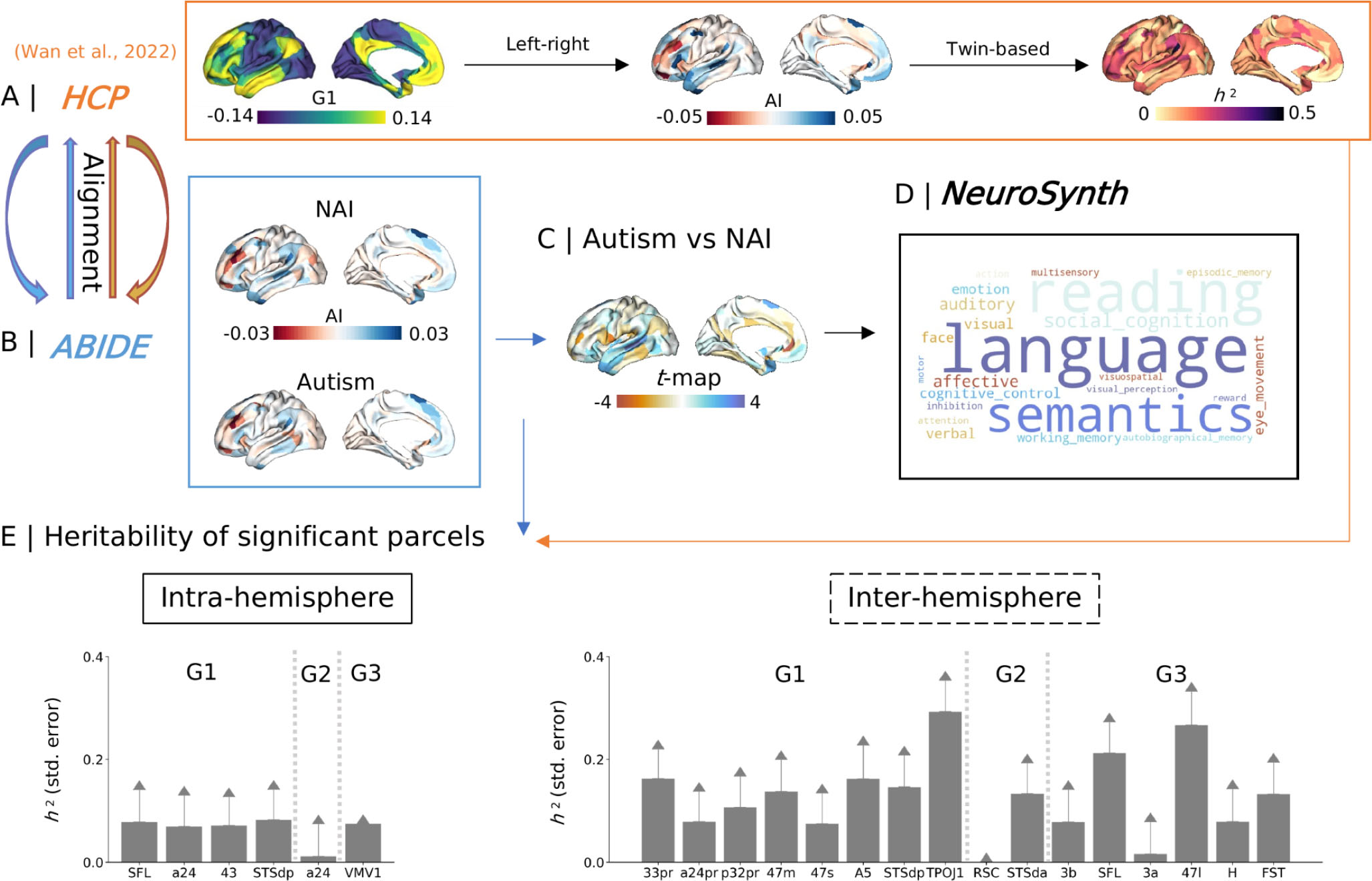
Heritability and meta-analytic decoding for *t*-map (G1). **A** | Yellow square shows how the heritability was computed by (Wan et al., 2022) using HCP twin-based data. Details can be seen in **Methods**. **B** | Blue square is the asymmetry in autism and NAI using the same pipeline to align individual gradients to the HCP template. **C** | Comparison between autism and NAI in asymmetry brain map. We showed the *t*-map here for functional decoding. **D** | Black square describes functional decoding of meta-analytic maps using NeuroSynth (Yarkoni et al., 2011). The bigger size of the text indicates the larger relevance to higher absolute *t*-values. The details regarding how to generate the maps and word clouds can be seen in the **Methods**. We also performed functional decoding for the other asymmetry maps (**Supplementary Results**). **E** | Illustrates the heritability with standard error estimates derived from panel A.

Regarding functional decoding, the *t*-map calculated from **Figure 3B** of intra-hemispheric G1 (**Figure 3C**) showed strong relevance to language, reading, and semantics (**Figure 3D**). The *t*-map of inter-hemispheric G1 showed strong relevance to language, reading, and semantics as well. Autism status*age effects along intra-hemispheric G3 showed strong relevance to auditory, language, and reading. Autism status*age effects along inter-hemispheric G3 showed strong relevance to language, auditory, and affective functions. Other functional decoding results are shown in **Supplementary results**.

Heritability is a marker that illustrates the proportion of variance across a population to be attributed to genetic factors. Here we sought to understand whether regions showing asymmetry differences between individuals with autism and NAI would be heritable within a population in young adulthood (22 - 37 years, HCP sample), as a proxy for a potential genetic versus environmental interplay associated with asymmetry. We extracted heritability values with standard error (SE) of the regions displaying diagnostic effects (**Figure 3E** and **Supplementary Table S7**). The regions displaying diagnostic effects along intra-hemispheric G1 showed low heritability, ranging from 0.069 to 0.083. The regions displaying diagnostic effects along inter-hemispheric G1 showed moderate heritability, ranging from 0.074 to 0.293, of which 33pr (*h*^2^ = 0.163, SE = 0.057, *p*_FDR_ = 0.006), 47m (*h*^2^ = 0.137, SE = 0.062, *p*_FDR_ = 0.027), A5 (*h*^2^ = 0.162, SE = 0.065, *p*_FDR_ = 0.015), STSdp (*h*^2^ = 0.146, SE = 0.062, *p*_FDR_ = 0.020), and TPOJ1 (*h*^2^ = 0.293, SE = 0.061, *p*_FDR_ < 0.001) survived after FDR correction. Moreover, STSda (*h*^2^ = 0.133, SE = 0.059, *p*_FDR_ = 0.038) along inter-hemispheric G2, SFL (*h*^2^ = 0.212, SE = 0.060, *p*_FDR_ = 0.001), 47I (*h*^2^ = 0.267, SE = 0.065, *p*_FDR_ < 0.001), and FST (*h*^2^ = 0.133, SE = 0.062, *p*_FDR_ = 0.028) along inter-hemispheric G3 survived after FDR correction.

### Phenotypic associations (Figure 4)

Lastly, we aimed to test whether asymmetry features (540 features based on 180 parcels * 3 gradients) can predict autistic traits as measured by ADOS (n = 132). To do so, we combined a linear regression with elastic net 5-fold cross validation (CV) with a supervised machine learning approach (**Figure 4A**) using scikit-learn (https://scikit-learn.org). Here we used L1_ratio = 0.1 to set up the regularization. Details using other L1_ratio parameters can be found in **Supplementary Results**.

**Figure 4.**
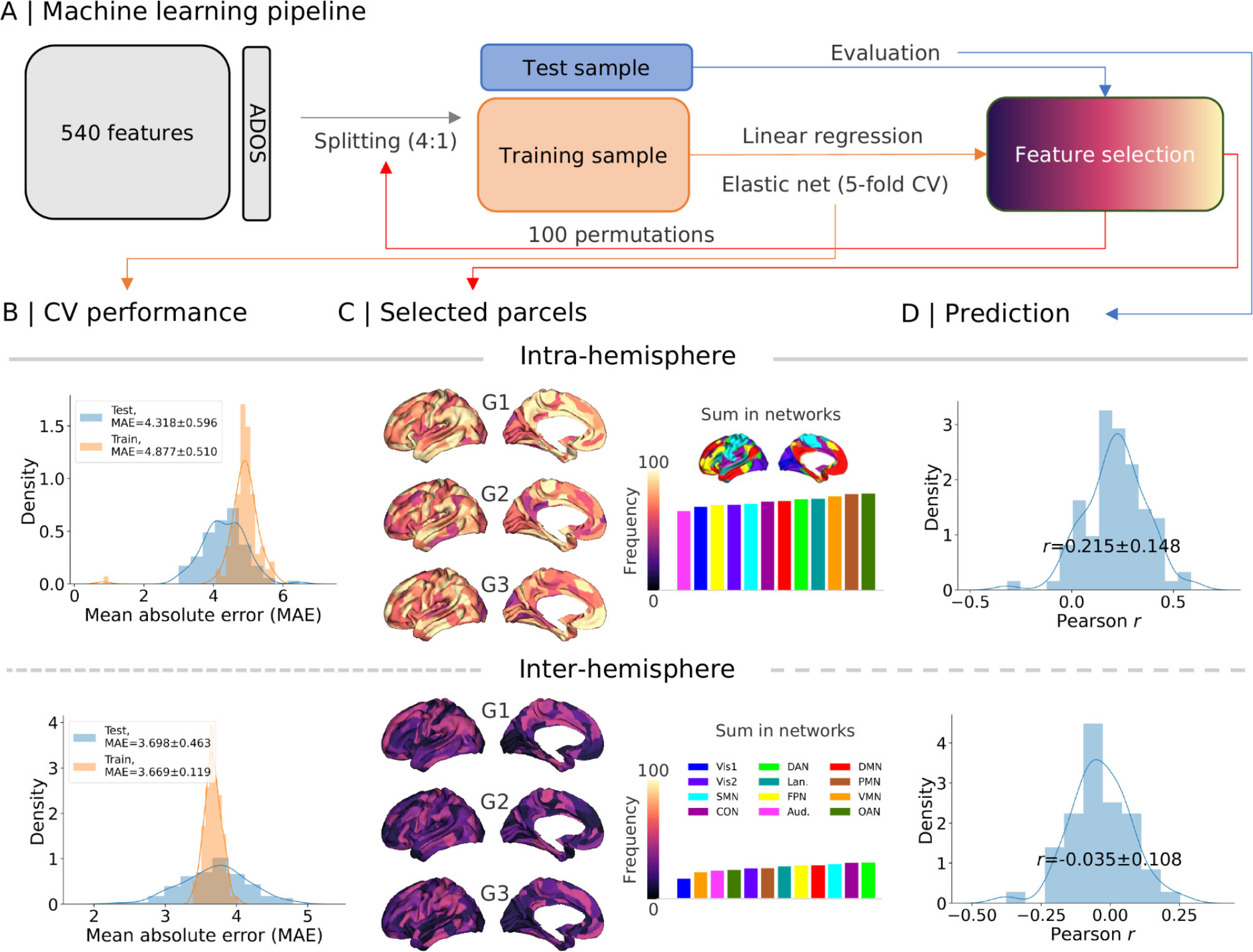
Autism traits and functional asymmetry. **A** | Machine learning pipeline. We used 540 features (180 parcels * 3 gradients) to predict the total score of ADOS. All subjects were split into 4:1 training:testing samples. Linear regression with elastic net (L1_ratio = 0.1) was the feature selector during which 5-fold cross validation was employed. We permuted this procedure 100 times by splitting the subjects randomly. **B |** Shows how the model works in the training and testing samples using mean absolute error (MAE) during the 100 permutations. **C** | Summarizes the frequency of this feature being selected across the 100 permutations. **D |** Illustrates the distribution of correlations between the observed ADOS total score and predicted ADOS total score in the testing samples across the 100 permutations. Vis1: primary visual network, Vis2: secondary visual network, SMN: somatomotor network, CON: cingulo-opercular network, DAN: dorsal attention network, Lan.:language network, FPN: frontoparietal network, Aud.: auditory network, DMN: default mode network, PMN: posterior multimodal network, VMN: ventral multimodal network, OAN: orbito-affective network.

Briefly, we randomized the whole sample for train-test (4:1) samples for 100 permutations. Multi-site and age effects were regressed out using neuroCombat data harmonization (Fortin et al., 2018) for training and testing samples separately. To automatically tune hyperparameters, we set a series of alphas from 0.0001 to 1. Elastic net with 5-fold CV estimated the models. The model with the averaged mean absolute error (MAE) was selected as being well-trained. Finally, to evaluate how much the model fit the testing sample, we calculated the Pearson *r* between the actual score and the predicted score in the testing sample. Across 100 permutations, MAE distributions are shown in **Figure 4B** with the mean ± SD being 4.875 ± 0.508 for the training sample and 4.314 ± 0.595 for the testing sample.

**Figure 4C** illustrates the frequency of each selected feature across the 100 permutations. In our pipeline, each feature had at least 50% possibility of being selected. High frequency occurred in DMN, PFN, and Lan. along G1, DAN, Lan., and OAN along G2, and Lan., FPN, and VMN along G3. Out-of-sample prediction suggests mean ± SD Pearson *r* to be 0.215 ± 0.148 for ADOS total score (**Figure 4D**). However, the models using inter-hemispheric asymmetry features were not good, with a mean ± SD Pearson *r* of -0.035 ± 0.108 for ADOS total score (**Supplementary Results**). Regarding prediction for ADOS subscores, intra-hemispheric asymmetry features fit positively for communication (Pearson *r =* 0.130 ± 0.142) and social (Pearson *r =* 0.153 ± 0.160) but not RRB (Pearson *r =* -0.097 ± 0.121), possibly in line with functional relevance of functional brain asymmetry.

## Discussion

In the current study, we examined the left-right asymmetry differences of functional connectome gradients between autism and NAI. Using a multivariate approach, we observed that sensory-motor and language networks showed diverging asymmetric organization in individuals with autism relative to NAI. In particular, the intrinsic functional asymmetry of language networks was related to altered developmental trajectories of functional organization, most prominently along the functional axis differentiating the default from multiple demand networks. Moreover, individuals with autism showed higher population heterogeneity for the spatial patterns of asymmetry. Intra-hemispheric asymmetry in language areas such as STS, found to differ between autism and NAI, were previously found to have low heritability in a young adult sample of NAI. However, group differences in inter-hemispheric asymmetry were observed to have moderate heritability strengths in NAI. These diverging patterns also suggest that both genetic and environmental components may be important to consider in the context of functional asymmetry differences between autism and NAI across development. Last, we found that, rather than inter-hemispheric asymmetry features, intra-hemispheric asymmetry features were more predictive of autistic traits. Together, our work shows extended differences in functional organization asymmetry between autism and NAI, which may be rooted in development and highly variable across individuals.

In the current work we leveraged the ABIDE sample, an openly available, multi-site cohort of individuals with autism and NAI (Di Martino et al., 2014, 2017) through data harmonization. Previous work on the same sample has revealed that both sensory and default regions are functionally more integrated in autism relative to NAI (Hong et al., 2019). Here, we extend these findings by showing that the language network had higher integration in the left hemisphere in NAI whereas in autism higher integration was found in the right hemisphere. Interestingly, networks that show rightward asymmetry in healthy individuals, e.g., FPN and CON along the sensory-default axis (Wan et al., 2022), are more rightward in individuals with autism compared with NAI. This differing processing pattern between hemispheres in autism is consistent with the idea of pervasive rightward lateralization in the disorder (Cardinale et al., 2013; Floris et al., 2013, 2016). Cardinale and colleagues (2013) used independent component analysis (ICA) to reveal 10/17 asymmetric networks and found that these networks (visual, auditory, motor, executive, language, and attentional) without exception display atypical rightward asymmetry in autism. Atypical motor performance in autism is correlated with their rightward motor circuits (Floris et al., 2016). These findings are mirrored by task-based reports, including language (Kleinhans et al., 2008; Knaus et al., 2008) and face processing (Solomon-Harris, 2022) tasks and may be linked to increased rightward or decreased leftward functional activation in autism (Kleinhans et al., 2008). As a system-level measurement, the gradients approach describes a regional feature as a function of an interregional embedding (Bernhardt et al., 2022). Thus, observed local regional asymmetries reported in prior work may result from systemic alterations in integration and segregation of functional connectivity as reported here.

Both cortical asymmetry and organization of intrinsic function show developmental change. For example, though left-right asymmetry is observed in the neonatal brain, frontal and temporal asymmetry in neonates differs from observations in adults (Williams et al., 2023). Here, we revealed a developmental component to deviating asymmetry of functional organization in language-related regions. Such a developmental alteration may be in line with reported delays in language and communication in autism (Luyster et al., 2007). The developmental trajectory of language-task activation lateralization follows an upwards trend from early childhood to adolescence, plateaus between 20 and 25 years, and slowly decreases between 25 and 70 years (Szaflarski et al., 2006). This converges with our observations in language network asymmetry using a system-level approach along G1, i.e., increased leftward asymmetry from childhood to adolescence and slightly decreased asymmetry from adolescence to adulthood in NAI. At the same time, we did not observe such developmental changes in autism, nor an interaction between autism status and age. This may indicate that the embedding of the language network, relative to attention networks, shows differential changes during development in individuals with autism relative to NAI, whereas its embedding between perceptual and abstract cognitive functions varies within autism, relative to NAI, irrespective of age. It has been suggested that initially bilateral language activation becomes more left-lateralized in typically developing children, whereas children with autism show a different developmental trajectory becoming increasingly rightward lateralized (Flagg et al., 2005). This indicates there should be an interaction with respect to language network asymmetry. We indeed observed such an interaction along G3. Language network asymmetry along G3 alters its direction from leftward to rightward during typical maturation whereas for autism we observed a subtle and leftward trend. It may suggest that language functions have multidimensional maturation trajectories (i.e., G1 and G3). The network-wise results were driven by language-related parcels in the temporal gyrus instead of frontal gyrus. It is of note that the current sample includes largely high-functioning individuals with autism. Yet, oral language impairments are observed in various degrees along the autism spectrum (Luyster et al., 2007) and language comprehension (especially under social context), instead of oral language, might be more apparent in individuals with high-functioning autism. Thus, in future work it will be relevant to study asymmetry in brain organization in a possibly more heterogeneous sample of individuals with autism.

Importantly, we observed age differences in functional gradient asymmetry in NAI but not in autism. This suggests that whilst asymmetry of functional organization in NAI changes over the course of development, this is not the case in individuals with autism. Other work also supports the notion that age effects of brain asymmetry are not found in individuals with autism (Guadalupe et al., 2017; Kong et al., 2018; Szaflarski et al., 2006; Zhou et al., 2013). This may reflect a maturation failure model in neurodevelopmental conditions and disorders (Di Martino et al., 2014), which in the case of autism may lead to different asymmetry development. Research shows that cortical asymmetries may largely be determined prenatally and that they may constrain the development of lateralized functions in later life (Williams et al., 2023). This suggests asymmetry is determined by genetics and environment *in utero*. In particular, environmental effects over the left hemisphere may be stronger than the right hemisphere *in utero* (Geschwind et al., 2002). Thus, the maturation alterations of brain asymmetry in autism might result from a complex interplay between genetic and environmental effects. Our study analyzed cross-sectional development, yet longitudinal data are necessary to evaluate the maturation failure model in autism.

Further investigating the interplay of developmental effects from genes on brain asymmetry, we evaluated whether regions showing differential asymmetry in autism are heritable in a normative non-autistic adult sample (Wan et al., 2022). We found that temporal language regions such as posterior STS and TPOJ1 along G1 between autism and NAI were heritable, whereas they were not heritable under intra-hemispheric connections. This different heritability of asymmetry patterns suggests the global feature in superior STS is more variable during intra-hemispheric specialization and may show stronger genetic constraints during inter-hemispheric specialization. Recent work using single-nucleotide polymorphisms (SNPs)-based analyses in the UK Biobank suggest high heritability in surface area asymmetry in these two regions. Further work, using more refined genetic imaging analysis, may help to further understand the neurobiological mechanisms underlying regional asymmetry and its functional consequences (Sha et al., 2021). Thus, inter-but not intra-hemispheric connectivity might be linked in some manner to additive genetic factors. As mentioned previously, environmental factors may have double effects on the left vs right hemisphere in brain volume *in utero* development (Geschwind et al., 2002). One possibility is that genes influence spatial organization in the right hemisphere, relevant to the inter-hemispheric function of superior STS, but that genes associated with autism impact inter-hemispheric connectivity. Further work using multilevel genetic and imaging data as well as brain models may help provide answers to these questions.

Lastly, we identified multiple areas related to autism traits via machine learning procedures similar to previous work (Hong et al., 2019; B. Park et al., 2021). This indicates that the model optimizes the parameters by averaging the features’ effects in autism and may reflect the complexity of autism traits using asymmetry features. We observed that the prediction using inter-hemispheric features is not as good as using intra-hemispheric features to predict ADOS total score. Follow-up indicated that subscores of communication and social traits showed acceptable out of sample prediction. Yet, repeated behaviors did not, underscoring the social and communicative trait relevance of individual variation in functional asymmetry. The differentiation between intra- and inter-hemispheric differences in terms of their predictability may again point to differential association between developmental and baseline effects. Indeed, intra- and inter-hemispheric asymmetry is primarily differentiated by the developmental timing of the role of corpus callosum (Gazzaniga, 2000; Toga & Thompson, 2003). Some studies have reported atypical cross-hemispheric connectivity and reduced corpus callosum size in autism compared to NTC (Anderson et al., 2011; Freitag et al., 2009). However, agenesis of the corpus callosum is not enough to specify the autistic traits (Paul et al., 2014). Future work may investigate the interplay between genes and environment in the context of intra- and inter-hemispheric connectivity and its clinical relevance to autism. Such differences may have already led to the differential idiosyncrasy of inter- and intra-hemispheric patterning, with idiosyncrasy of inter-hemispheric connectivity to be more extended in autism.

Overall, through studying the organization of intrinsic functional asymmetry, our work provides a framework to study hemispheric differences in individuals with autism versus NAI. However, there are several limitations to note. First of all, the current study was based on neuroimaging data from multiple acquisition sites, enhancing the sample size but at the same time also introducing potential site-related confounds. We used data harmonization (Fortin et al., 2018) to reduce this influence as much as possible. Second, the enrichment decoding results are indirect. If we want to understand the genetic basis and cognitive relevance of asymmetry in autism and healthy individuals, it is necessary to measure genetic and cognitive features in autism. Moverover, in the current sample, we could not provide the causal link between development and functional asymmetry. Longitudinal design and/or high-risk autism models may help to highlight the neurodevelopmental foundations of functional asymmetry in autism and guide implications for support. Finally, we excluded autistic females, however, research shows that brain lateralization differs by sex (Kong et al., 2018; Liang et al., 2021; McGlone, 1978, 1980) and autism shows sex and gender differences in prevalence, behavior and brain (Floris et al., 2023; Fombonne, 2003; Hull et al., 2017; Lai et al., 2015; Xu et al., 2018). Future studies should investigate whether there exist sex and gender-differential patterns of atypical asymmetry in autism.

To conclude, we report functional organization asymmetry in autism, its age-related changes, and trait relevance. In particular, we detected decreased leftward asymmetry in the language network along the sensory-default gradient and somatomotor network along the default-multiple demand gradient in autism. A differing developmental trajectory in autism was observed in the language network along the default mode-multiple demand gradient. Moreover, functional asymmetry is a central feature of autism, linking to autistic traits, with marked deviations from controls in terms of development and idiosyncrasy. Future work may study the impact of environmental factors upon genes associated with autism during early development and associated traits and cognitive development across the lifespan.

## Methods

We employed five datasets that covered children, adolescents, and young adults from the Autism Brain Imaging Data Exchange (ABIDE, http://fcon_1000.projects.nitrc.org/indi/abide), of which ABIDE-I includes New York University Langone Medical Center (NYU-I), University of Pittsburgh-School of Medicine (Pitt), and University of Utah-School of Medicine (USM), and ABIDE-II includes NYU-II and Trinity Centre for Health Sciences-Trinity College Dublin (TCD). In accordance with HIPAA guidelines and 1000 Functional Connectomes Project / INDI protocols, all ABIDE datasets have been anonymized, with no protected health information included.

### Participants

We restricted our analyses to males (n = 300) due to the low number of females with autism, consistent with previous work (Hong et al., 2019). Individuals with autism underwent a structured or unstructured in-person interview and had a diagnosis of Autistic, Asperger’s, or Pervasive Developmental Disorder Not-Otherwise-Specified. These were established by expert clinical opinion aided by ‘gold standard’ diagnostics: Autism Diagnostic Observation Schedule Generic version (ADOS-G, Lord et al., 2000) and/or Autism Diagnostic Interview-Revised (ADI-R). Subdomains include communication, social interaction, and restricted repetitive behaviors (RRB). Intelligence quotient (IQ) was measured by the Wechsler Abbreviated Scale of Intelligence including III, IV, and V versions (Wechsler, 1999).

We excluded subjects with age greater than 40 years (n = 2) to retain a centralized population age and full IQ below 70 (n = 1) to avoid developmental delay of intelligence. Regarding head motion, we measured mean framewise displacement (FD), derived from Jenkinson’s relative root mean square algorithm (Jenkinson et al., 2002). We excluded individuals whose mean FD was greater than 0.3 mm (n = 14), consistent with the previous report (Hong et al., 2019). The final sample size taken into analyses was n = 283. Among these, we categorized them into three age groups including 142 young adults (18 - 40 years, autism: n = 66), 97 adolescents (12 - 17 years, autism: n = 51), and 76 children (6 - 11 years, autism: n = 40).

To reduce the effects of data sites, we conducted data harmonization (Fortin et al., 2018) using the toolbox neuroCombat (https://github.com/Jfortin1/neuroCombat). It provides a Bayesian approach to balance the effects of each scanner/site as well as continuous or categorized covariates. FIQ and head motion were entered as covariates during data harmonization. Results remain consistent and can be seen in our online iPython notebook.

### Preprocessing of resting state fMRI data

High-resolution T1-weighted images (T1w) and resting-state functional magnetic resonance imaging (fMRI) data were available from all five sites. The scanning parameters and preprocessing procedures are reported in previous work (Hong et al., 2019). In short, 3D-TurboFLASH was used for T1w of NYU datasets and 3D-MPRAGE was used for T1w of the other three datasets. TR ranged from 2100 to 300 ms and TE from 2.91 to 3.90 ms. The resolution was 1.1*1.0*1.1 mm^3^ voxels. A 2D EPI sequence was employed for resting state fMRI data with the TR ranging from 1500 ms to 2000ms, volumes ranging from 180 to 240, and a resolution of 3.0*3.0*3.4 mm^3^ voxels.

T1w data processing was done with FreeSurfer (v5.1; http://surfer.nmr. mgh.harvard.edu/). Image processing included bias field correction, registration to stereotaxic space, intensity normalization, skull-stripping, and white matter segmentation. Our fMRI analysis was based on preprocessed data previously made available by the Preprocessed Connectomes initiative (http://preprocessed-connectomesproject.org/abide/). Preprocessing was based on C-PAC (https://fcp-indi.github.io/) and included slice-time correction, head motion correction, skull stripping, and intensity normalization. Statistical corrections removed effects of head motion, white matter, and cerebrospinal fluid signals using the CompCor tool, based on the top 5 principal components, as well as linear/quadratic trends. After band-pass filtering (0.01 - 0.1 Hz), we co-registered resting state fMRI and T1w data in MNI152 space through combined linear and non-linear transformations.

Surface alignment was verified for each case and we interpolated voxel-wise rs-fMRI time-series along the mid-thickness surface. We resampled rs-fMRI surface data to downsampled Conte69 (10k vertices per hemisphere), a template mesh from the Human Connectome Project pipeline (https://github.com/Washington-University/Pipelines), and applied surface-based smoothing (FWHM = 5 mm). MRI quality control was complemented by assessment of signal-to-noise ratio and visual scoring of surface extractions for T1w.

### Parcellation

To reduce the high computational demands of processing vertex-based fMRI data, we downsampled vertex-based fMRI data to 180 parcels per hemisphere using multimodal parcellation (MMP, Glasser et al., 2016) and summarized features into Cole-Anticevic (CA) 12 functional networks (Ji et al., 2019). MMP has been generated using the gradient-based parcellation approach with similar gradient ridges presenting in roughly corresponding locations in both hemispheres, which is suitable for studying asymmetry across homologous regions. Regarding cortical functional communities, CA atlas summarizes 12 functional networks based on MMP including primary visual (Vis1), secondary visual (Vis2), somatosensory (SMN), cingulate-opercular (CON), dorsal attention (DAN), language (Lan.), frontoparietal (FPN), auditory (Aud.), default mode (DMN), posterior-multimodal (PMN), ventral-multimodal (VMN), and orbito-affective (OAN).

### Functional connectome gradients

After parcellating the preprocessed signals, we can get the arrays of time series * parcels. We first computed the Pearson correlation between parcels using time series and transformed r values to z values using Fisher z-transformation. This generates the functional connectivity (FC) matrices of 360*360 for each individual. Then, to compute the functional connectome gradients, we used a non-linear manifold learning algorithm, to perform dimensionality reduction of the FC matrix. Consistent with the framework of asymmetry of functional gradients (Wan et al., 2022), we aligned each individual gradient to the template gradient (i.e., left-left group level gradients) with Procrustes rotation to make individual gradients comparable (VosdeWael, 2020). To gain an unbiased left-left group-level gradients template, without age or gender bias, in young adults, we employed data from Human Functional Connectome project S1200 release (HCP S1200). This has been done previously (Wan et al., 2022). Briefly, we averaged 1104 subjects FC matrices of HCP S1200 and computed the group level gradients based on the mean left-left FC matrix. The first eigenvectors reflect unimodal-transmodal gradient (G1), sensory-visual gradient (G2), and multi-demand gradient (G3) explaining 24.1, 18.4, and 15.1% of total variance each.

Gradient analysis was performed in BrainSpace (Vos de Wael et al., 2020), a Matlab/python toolbox for brain dimensionality reduction (https://brainspace.readthedocs.io/en/latest/pages/install.html). Gradients are low dimensional eigenvectors of the connectome, along which cortical nodes that are strongly interconnected, by either many suprathreshold edges or few very strong edges, are situated closer together. Similarly, nodes with little connectivity are farther apart. This reflects the integration and segregation between regions located on the common spaces. The name of this approach, which belongs to the family of graph Laplacians, is derived from the equivalence of the Euclidean distance between points in the diffusion map embedding (Coifman & Lafon, 2006; Margulies et al., 2016). It is controlled by a single parameter α, which reflects the influence of the density of sampling points on the manifold (α=0, maximal influence; α=1, no influence). On the basis of the previous work (Margulies et al., 2016), we followed recommendations and set α = 0.5, a choice that retains the global relations between data points in the embedded space and has been suggested to be relatively robust to noise in the covariance matrix. The top 10% of values in the FC matrix were used for the threshold to enter the computation, consistent with previous studies (Hong et al., 2019; Margulies et al., 2016; Wan et al., 2022).

### Asymmetry index

To quantify the left and right hemisphere differences, we chose left-right as the asymmetry index (AI) (Wan et al., 2022). We did not opt for normalized AI, i.e., (left-right)/(left +right), as gradient variance (normalized angle) has both negative and positive values (Sha et al., 2022) and normalized AI exaggerates the difference values or results in a discontinuity in the denominator (Nielsen et al., 2013). The normalized AI is highly similar to non-normalised AI with correlation coefficients greater than 0.9 (Wan et al., 2022). For the intra-hemispheric pattern, the AI was calculated using LL-RR. A positive AI-score meant that the hemispheric feature dominated leftwards, while a negative AI-score dominated rightwards. For the inter-hemispheric pattern we used LR-RL to calculate the AI. We added a ‘minus’ to Cohen’s d scores in the figures in order to conveniently view the lateralization direction (i.e., leftward or rightward).

### Heritability and meta-analytic decoding

Regarding the meta-analytic decoding, we used activation data from NeuroSynth database (Yarkoni et al., 2011). We chose 24 cognitive domain terms, consistent with previous studies (Margulies et al., 2016; Wan et al., 2022). In the present study, to decode both hemispheres, *t*-values of leftward AI were directly put on the right hemisphere and *t*-values * 2 of leftward AI were put on the left hemisphere. Similarly, *t*-values of rightward AI were put on the left hemisphere and *t*-values * 2 of rightward AI were put on the right hemisphere. We generated 20 bins for the brain map averagely (5% per bin). Thus, each cognitive domain term had a mean activation z-score per bin. To assess how much the cognitive domain terms are abnormal in autism, we calculated a weighted score by mean activation (where activation z-score greater than 0.5) multiplied by *t*-values. A bigger shape in the word cloud reflects a higher weighted score (i.e., atypical lateralized functions in autism).

The heritability data were derived from a prior study by our team (Wan et al., 2022) that was based on a study of non-autistic adult twins/non-twins. After selecting the parcels where autism showed differences from NAI, we could describe their genetic underpinnings with heritability data from HCP.

### Prediction

We performed supervised machine learning to predict the ADOS total and subscale scores. Regarding cross-validation, we applied a 5-fold leave-one-out strategy to learn the data. Among the 5-time iterations, the one with averaged mean absolute error (MAE) was chosen as the final model to predict the clinical symptoms. Linear regression with elastic net (L1_ratio = 0.1) was used as the feature selector. This follows an Empirical Bayesian approach to balance the effects of each scanner/batch. After the features’ contributions had been built, we used Pearson correlation coefficients to evaluate how strong the model could be applied to the current sample.

First, we divided the participants into training and testing samples using a 4 to 1 ratio. Next, we applied data harmonization for training and testing samples separately. We then used the cross-validation as described above to select features in the training sample. The selected features were then fit in the independent testing sample to evaluate the model. We permuted the whole procedure 100 times with a random number to split the subjects into training and testing samples. This enabled us to know the frequency of how often features are selected over the 100 permutations.

### Data and code availability

The ABIDE open data can be acquired from https://fcon_1000.projects.nitrc.org/indi/abide/. All the analysis scripts and visualization for this study are openly available at a Github repository (https://github.com/wanb-psych/autism_gradient_asymm). Key dependencies are Python 3.9 (https://www.python.org/), BrainSpace (https://brainspace.readthedocs.io/), and BrainStat (https://brainstat.readthedocs.io/).

## Acknowledgements

We are grateful to the open access platform for neuroimaging data sharing in autism: ABIDE (https://fcon_1000.projects.nitrc.org/indi/abide/). B.C.B. acknowledges research support from the National Science and Engineering Research Council of Canada (NSERC Discovery-1304413), Canadian Institutes of Health Research (FDN-154298, PJT-174995), SickKids Foundation (NI17-039), BrainCanada, FRQ-S, and the Tier-2 Canada Research Chairs program. S.L.V and B.C.B are furthermore funded by the Helmholtz International BigBrain Analytics and Learning Laboratory (HIBALL), supported by the Helmholtz Association’s Initiative and Networking Fund and the Healthy Brains, Healthy Lives initiative at McGill University. S.L.V is supported by the Otto Hahn Award at Max Planck Society, and B.W is supported by the International Max Planck Research School on Neuroscience of Communication: Function, Structure, and Plasticity (IMPRS NeuroCom).

## Notes

### Competing Interest Statement

The authors have declared no competing interest.

